# Expression and distribution of Trophoblast Glycoprotein in the mouse retina

**DOI:** 10.1101/590216

**Authors:** Colin M Wakeham, Gaoying Ren, Catherine W Morgans

## Abstract

We recently identified the leucine-rich repeat adhesion protein, trophoblast glycoprotein (TPBG), as a novel PKCα-dependent phosphoprotein in retinal rod bipolar cells (RBCs). Since TPBG has not been thoroughly examined in the retina, this study characterizes the localization and expression patterns of TPBG in the developing and adult mouse retina using two antibodies, one against the N-terminal, leucine-rich domain and the other against the C-terminal PDZ-interacting motif. Both antibodies labeled dendrites and synaptic terminals of RBCs, as well as the cell bodies and dendrites of an uncharacterized class of amacrine cell. In transfected HEK293 cells, TPBG was localized to the plasma membrane and intracellular membranes and was associated with the tips of thin filopodia-like membrane projections. TPBG immunofluorescence in RBCs detected with the C-terminal antibody was strongly dependent on the activity state of the adult retina, with less labeling in dark-adapted compared to light-adapted retina, and less labeling in light-adapted PKCα knockout and TRPM1 knockout retinas compared to wild type, despite no change in total TPBG detected by immunoblotting. These results suggest that the C-terminal epitope is blocked in the dark-adapted and knockout retinas compared to light-adapted wild type retinas, possibly through interaction with a PDZ domain protein. During development, TPBG expression increases dramatically just prior to eye opening with a time course closely correlated with that of TRPM1 expression. In the retina, leucine-rich repeat proteins like TPBG have been implicated in the development and maintenance of functional bipolar cell synapses, and TPBG may play a similar role in RBCs.

## 1. Introduction

Rod bipolar cells (RBCs) are the first excitatory interneurons in the rod visual system. They receive light-dependent synaptic input from rod photoreceptors in the outer plexiform layer (OPL) and contribute to retinal output via AII amacrine cells in the inner plexiform layer (IPL). RBCs are associated with dark-adapted vision (Euler, Haverkamp, Schubert, & Baden, 2014), yet recent evidence suggests that RBCs contribute to retinal output under a diverse range of lighting conditions. Under completely dark-adapted conditions, RBCs are sensitive to single-photon responses in rods (Berntson, Smith, & Taylor, 2004; Sampath & Rieke, 2004). In mesopic conditions, RBCs contribute to the perception of contrast (Abd-El-Barr et al., 2009; Ke et al., 2014). Finally, in bright light, RBCs may modulate the cone pathway when rods are saturated (Szikra et al., 2014). The molecular mechanisms required for RBC adaptation to changing luminance conditions are mostly unknown, but compelling evidence implicates the commonly-used RBC marker protein kinase C-alpha (PKCα; Rampino & Nawy, 2011; Ruether et al., 2010; Xiong et al., 2015).

To gain insight into the mechanisms by which PKCα modulates the RBC light response, we sought to identify RBC proteins that undergo PKCα-dependent phosphorylation. Using a multiplexed tandem mass tag mass spectroscopy-based approach, we previously identified trophoblast glycoprotein (TPBG, also known as 5T4 or WAIF1 [Wnt-activated inhibitory factor 1]) as a novel PKCα-dependent phosphoprotein in RBCs (Wakeham et al., 2019). TPBG is a type 1 transmembrane glycoprotein with an N-terminal extracellular domain composed of eight leucine rich repeats interspersed by seven N-linked glycosylation sites. The intracellular cytoplasmic domain is capped by a class 1 PDZ-interacting motif (Zhao, Malinauskas, Harlos, & Jones, 2014) and contains two serines which were significantly more likely to be phosphorylated in wild type retinas compared to PKCα knockout (Wakeham et al., 2019).

TPBG was first identified in trophoblasts (Hole & Stern, 1988) and has been mainly studied in embryonic development and in cancer (Barrow, Ward, Rutter, Ali, & Stern, 2005), where it is required for cell surface expression of CXCR4, a G-protein coupled receptor that mediates a chemotactic response to the CXCL12 chemokine (McGinn, Marinov, Sawan, & Stern, 2012; Southgate et al., 2010), and where it is diagnostic for metastasis and poor prognosis in cancer patients (Pukrop & Binder, 2008; Weeraratna et al., 2002). In mammalian embryonic cell lines, TPBG influences cytoskeletal organization and cell motility through modulation of Wnt signaling (Kagermeier-Schenk et al., 2011), and has also shown to bind to the PDZ domain of GIPC1, a scaffolding protein that regulates cell surface expression of G protein-coupled receptors (Awan et al., 2002). In adult tissues, it is expressed at high levels in ovary, brain, and retina (Imamura et al., 2006; King, Sheppard, Westwater, Stern, & Myers, 1999). Relatively little is known about the role of TPBG in neurons; however, in the olfactory bulb, TPBG has been shown to drive developmental changes in the dendritic morphology of granule cells in an activity-dependent manner, and genetic knockdown of TPBG resulted in impaired odor discrimination (Takahashi et al., 2016; Yoshihara et al., 2012; Yoshihara, Takahashi, & Tsuboi, 2016).

In the retina, a comprehensive single-cell drop-seq study identified TPBG as a possible new RBC marker and clustered it with proteins involved in lamination and adhesion (Shekhar et al., 2016). In a subsequent comparative meta-analysis of RBC transcriptomics studies, it was grouped as a protein potentially involved in synapse assembly (Woods, Mountjoy, Muir, Ross, & Atan, 2018). As TPBG has never been thoroughly examined in RBCs, we characterized its expression and subcellular localization patterns in both the adult and developing mouse retina.

## 2. Methods

### Antibody characterization

Rabbit anti-5T4 (TPBG-CT) (1:500 [immunofluorescence], 1:5000 [immunoblot]; Abcam; Cambridge, UK; Cat# ab129058; RRID: AB_11144484) is a monoclonal antibody targeting a 15-amino acid synthetic peptide immunogen corresponding to a sequence within amino acids 385-420 of human TPBG. TPBG immunoreactivity was verified by the manufacturer via immunoblot of rat brain, HeLa cell, and HT-1376 cell lysates, and immunoprecipitation from MCF-7 cell lysates and detected broad bands of glycosylated proteins at 70-80 kDa (manufacturer information).

Sheep anti-5T4 (TPBG-NT) (1:500; R&D Systems; Minneapolis, MN, USA; Cat# AF5049; RRID: AB_2272148) is a polyclonal antibody raised against amino acids 30 to 361 of mouse TPBG. The manufacturer used flow cytometry and immunocytochemistry to detect TPBG in the membranes of retinoic acid-treated D3 mouse cell lines (manufacturer information). We confirmed specificity of both the TPBG-CT and TPBG-NT antibodies by immunoprecipitation followed by immunoblotting from while type mouse retinal lysate and TPBG-transfected HEK293 cells, with both antibodies detecting a broad band around 72 kDa.

Mouse anti-PKCα (1:5000; Sigma-Aldrich; St. Louis, MO, USA; Cat# P5704; RRID: AB_477375) is a monoclonal antibody that recognizes an epitope between amino acids 296-317 on the hinge region of mouse PKCα. The manufacturer detected bands at 80 kDa in immunblots of lysates from SH-SY5Y, SK-N-SH, COS7, and PC12 cell lines (manufacturer information), and we have verified this antibody by immunoblots of retinal lysates from wild type and PKCα-KO mice.

Mouse anti-CANA1S (1:4000; anti-GPR179; Abcam; Cat# ab2862; RRID: AB_2069567) is a monoclonal antibody raised against full-length native rabbit CACNA1S subunit purified from the rabbit muscle T-tubule dihydropyridine receptor. This CACNA1S antibody has been shown to strongly cross-react with retinal GPR179 in the tips of ON-bipolar cell dendrites (Hasan, Ray, & Gregg, 2016).

Mouse anti-CtBP2 (1:4000; BD Biosciences; San Jose, CA, USA; Cat# 612044; RRID: AB_399431) is a monoclonal antibody that binds an epitope within amino acids 361-445 of mouse CtBP2. The manufacturer used western blot analysis on BC3H1 cell lysates and detected a band at 48 kDa (manufacturer information). This antibody produces strong immunoreactivity in both nuclei and horseshoe-shaped synaptic ribbons in wild type mouse retina sections, as is characteristic of CtBP2 (Schmitz, Königstorfer, & Südhof, 2000).

Mouse anti-calretinin (1:25; Santa Cruz Biotechnology; Dallas, TX, USA; Cat# sc-365956; RRID: AB_10846469) is a monoclonal antibody that recognizes an epitope between amino acids 2-27 at the N-terminus of human calretinin. The manufacturer used western blot analysis to detect bands between 23 and 34 kDa from human cerebellum, human brain, human adrenal gland, and rat cerebellum lysates (manufacturer information). We confirmed specificity by labeling wild type mouse retina sections and detected the characteristic three-band pattern of calretinin in the IPL.

Sheep anti-TRPM1 (1:2000) is a polyclonal antibody raised against a fragment of recombinant polypeptide corresponding to amino acids 1423-1622 of mouse TRPM1 (Y Cao, Posokhova, & Martemyanov, 2011). We received this antibody from Kirill Martemyanov.

Mouse anti-β-actin (1:2500; Cell Signaling Technology; Danvers, MA, USA; Cat# 8H10D10; RRID: AB_2242334) is a monoclonal antibody that was verified in this study by immunoblot of wild type whole retinal lysates from mice at different developmental stages and shown to produce bands at ∼42 kDa. The manufacturer used western blot analysis of extracts from COS, HeLa, C2C12, C6, and CHO cells to confirm immunoreactivity for β-actin.

The secondary antibodies used were AF488 anti-rabbit (1:1000; Jackson ImmunoResearch Labs; West Grove, PA, USA; Cat# 11-545-144; RRID: AB_2338052), AF488 anti-sheep (1:1000; Jackson ImmunoResearch Labs; Cat# 713-545-147; RRID: AB_2340745), Cy3 anti-mouse (1:1000; Jackson ImmunoResearch Labs; Cat# 115-165-003; RRID: AB_2338680), 680RD anti-rabbit (1:15000; LI-COR Biosciences; Lincoln, NE, USA; Cat# 925-68071; RRID: AB_2721181), 680RD anti-goat (sheep) (1:15000; LI-COR Biosciences; Cat# 925-68074; RRID: AB_265-427), and 800CW anti-mouse (1:15000; LI-COR Biosciences; Cat# 925-32212; RRID: AB_2716622).

### Expression vector

The TPBG expression vector contains full-length mouse TPBG cDNA (NM_011627.4) inserted into the pCMV-sport6 plasmid (Thermo Scientific; Cat# 12209) provided by the PlasmID Repository (Clone ID: MmCD00318800; Species ID: 21983; Harvard Medical School; Boston, MA, USA).

### Mice

Wild type mice used were C57BL/6J (Jackson Laboratory; Bar Harbor, ME, USA; Cat#000664, RRID: IMSR_JAX:000664). The PKCα knockout mice were B6;129-Pkrca^tm1Jmk^/J (Jackson Laboratory; Cat# 009068; Xiong et al., 2015). The TRPM1 knockout mice were TRPM1^tm1Lex^ (Texas A&M Institute of Genomic Medicine; College Station, TX, USA; Morgans et al., 2009). Mice of both sexes were used, and all mice were maintained on a 12-hour light/dark cycle and provided food and water ad libitum. Mice used for dark adaption experiments were maintained in darkness for 12 hours preceding tissue collection. All animal procedures were in accordance with the National Institutes of Health guidelines and approved by the Oregon Health and Science University Institutional Animal Care and Use Committee.

### HEK293 cell transfection

HEK293 cells (ATCC Cat# CRL-1573) were maintained at 37° C and 5% CO_2_ in 1X DMEM medium (Thermo Scientific; Waltham, MA, USA; Cat# 11995-065, RRID: CVCL_0045) supplemented with 10% fetal bovine serum (Gemini Bio-Products Cat# 900-108) and 1% Pen/Strep (Thermo Scientific; Cat# 15140-122). For protein expression, approximately 10^5^ cells were plated on coverslips (Thermo Scientific; Cat# 12-540-80) coated with poly-l-lysine (Sigma-Aldrich; Cat# P4704) in a 24-well dish. The next day, 0.2 ng of the pCMV-TPBG-sport6 vector was transfected using the Effectene transfection reagent kit (Qiagen; Venlo, Netherlands; Cat# 301525), and expression was assessed by immunostaining approximately 24 hours after transfection.

### Immunostaining transfected HEK293 cells

#### Fixed and permeabilized cells

Cells grown on glass cloverslips were washed with 0.1 M +Mg^2+^/+Ca^2+^ phosphate buffered saline, pH 7.4 (PBS), fixed for 10 min in 4% paraformaldehyde (PFA), and then washed again with PBS. The cells were permeabilized and blocked by incubation with antibody incubation solution (AIS: 3% normal horse serum, 0.5% Triton X-100, 0.025% NaN_3_ in PBS) for 30-60 min. Primary antibody diluted in AIS was added to the cells and incubated at room temperature for 1 hr, before being removed and the cells washed with PBS. Secondary antibody, also diluted in AIS, was added at room temperature for 1 hr, then removed. 1X DAPI was added to the coverslips for 1 min, before being washed off with PBS. The coverslips were then mounted on Super-Frost glass slides in Lerner Aqua-Mount (Thermo Scientific; Cat# 13800) and sealed.

#### Live cells

Cells grown on coverslips were washed with PBS and given fresh DMEM medium. Primary antibody was added directly to the medium, and the cells were placed on ice for 1 hr, after which they were washed with PBS, fixed with 4% PFA, and permeabilized with AIS. The remaining steps were identical to the previous section.

### Tissue preparation for immunofluorescence

#### Adult mouse retina sections

Mouse eyecups were prepared from freshly dissected eyes by cutting behind the ora serrata and removing the cornea and lens. Eyecups were fixed for 30 min by immersion in 4% PFA in PBS. The fixed eyecups were washed in PBS and then cryoprotected via sequential immersion in 10, 20, and 30% sucrose in PBS. The tissue was embedded in Tissue-Tek O.C.T. Compound (Sakura Finetek; Tokyo, Japan; Cat# 4583) and stored frozen at −80° C until sectioning. Sections were cut at 20-25 μm thickness on a cryostat and then mounted onto Super-Frost glass slides. The slides were air dried and stored at −20° C.

#### Developing mouse retina sections

Mouse eyes were freshly dissected and a small cut was made behind the ora serrata to allow fixative and cryoprotectant penetration into the retina. The whole eye was then fixed for 30 min by immersion in 4% PFA in PBS. The fixed eyes were then cryoprotected via sequential immersion in 10, 20, and 30% sucrose in PBS until they sank. The remaining steps were identical to the previous section.

### Immunostaining retina sections

Retina sections were thawed and then blocked and permeabilized by incubation at room temperature for 30-60 min in AIS, and then were incubated in primary antibodies for 1 hr at room temperature. After washing with PBS, the sections were incubated in secondary antibodies diluted in AIS for 1 hr at room temperature. Finally, the sections were incubated for 1 min in 1X DAPI. The slides were washed again in PBS and then mounted with Lerner Aqua-Mount. For the TPBG-CT antibody, retina sections were post-fixed for 15 min in 4% PFA before the blocking and permeabilization step as this was found to improve the immunofluorescence with this antibody.

### Scanning confocal imaging

Immunofluorescence images were taken with a Leica TCS SP8 X white light laser confocal microscope (Leica; Wetzlar, Germany) using a Leica HC PL APO CS2 63x/1.40 oil immersion objective (Leica; Cat# 15506350) and Leica HyD hybrid detectors. Laser lines used were DAPI (405 nm), AF488 (499 nm), Cy3 (554 nm), and AF594 (598 nm). Detection windows used were DAPI (415-489nm), AF488 (509-544nm), Cy3 (564-758 nm), and AF594 (608-766nm). Z-projections intended for comparison were processed in LAS X using identical tissue thicknesses and a “Threshold” setting of 20. Brightness and contrast were adjusted equally across comparison groups using Leica LAS X or ImageJ (Rueden et al., 2017). The ImageJ “Smooth” tool was used to remove graininess from images. Figure 3C was deconvolved with the Huygens HyVolution 2 module (Scientific Volume Imaging; The Netherlands) with the “standard” algorithm and a refractive index of 1.454.

### Immunoblotting

Retinas were extracted from freshly dissected eyes, suspended in chilled lysis buffer (50 mM Tris pH 7.4, 150 mM NaCl, 1 mM EDTA, 1% Triton X-100, 1% deoxycholate, 0.1% SDS) with 1X protease/phosphatase inhibitor cocktail (Cell Signaling Technology; Danvers, MA, USA; Cat# 5872) and homogenized with a Teflon-glass homogenizer. The lysate was centrifuged for 15 min at 16,400 rpm and 4° C, and the pellet was discarded. Lysates were stored at −20° C. Retinal lysates were diluted to 1μg/μl in lysis buffer and brought to 1X NuPAGE LDS Sample Buffer (Thermo Scientific; Cat# NP0007) and 1X NuPAGE Sample Reducing Agent (Thermo Scientific; Cat# NP0009). Pre-cast NuPAGE 1mm 4-12% Bis-Tris gels (Thermo Scientific; Cat# NP0322BOX) were loaded and run at 200 V and 140 mA for 55 min in 1X NuPAGE BOLT SDS Running Buffer (Thermo Scientific; Cat# B0001). Proteins were transferred onto PVDF membranes using a semi-dry transfer system and 2X NuPAGE Transfer Buffer (Thermo Scientific; Cat# NP00061) with 10% MeOH at 45 mA for 2 hrs, or a wet transfer system and 1X NuPAGE Transfer Buffer with 5% MeOH at 300mA for 2 hrs. The membranes were then rinsed with methanol and blocked for 1 hr in Odyssey Blocking Buffer TBS (LI-COR Biosciences; Cat# 927-50003) on a shaker at room temperature, before being incubated in primary antibody diluted in Odyssey buffer at 4° C overnight. The membranes were washed 3×5 min in TBST (Tris-buffered saline with 0.1% Tween-20), then incubated in secondary antibody diluted in Odyssey buffer for 1 hr at room temperature before being washed 3×5 min in TBST and left to dry. The dry blots were imaged using a LI-COR Odyssey CLx Imaging System at 700 and 800 nm.

## 3. Results

### TPBG is expressed in the dendrites and synaptic terminals of rod bipolar cells

Two commercial antibodies against different epitopes of TPBG were purchased. The first, a rabbit monoclonal (TPBG-CT), reacts with an epitope near the PDZ-interacting motif of TPBG’s intracellular C-terminal tail, while the second, a sheep polyclonal (TPBG-NT), binds within the extracellular N-terminal leucine-rich repeat (LRR) domain of TPBG.

Wild type adult mouse retina sections were labeled with either anti-TPBG-CT or anti-TPBG-NT, and the nuclear stain DAPI was used to identify the different retinal layers. Immunofluorescent labeling with anti-TPBG-CT (Figure 1A) reveals strong immunoreactivity in the OPL and in two bands in the IPL, one in the middle of the IPL at sublamina 2/3 and the other at the innermost IPL in sublamina 5. In the inner nuclear layer, anti-TPBG-CT labels putative RBC cell bodies as well as a sparse population of amacrine cell bodies. In a dissociated cell preparation (Figure 1B), TPBG-CT only labeled PKCα-positive RBCs. Immunofluorescent labeling with anti-TPBG-NT (Figure 1C) reveals punctate immunoreactivity in the OPL and widespread, punctate labeling throughout the IPL, with the strongest IPL labeling occurring in sublamina 5. In the INL, anti-TPBG-NT faintly labeled occasional amacrine cell bodies, but labeling of bipolar cell bodies was not observed. Although anti-TPBG-NT did not label bipolar cell bodies in retina sections, it clearly labeled PKCα-positive RBCs in a dissociated retina preparation (Figure 1D).

**Figure 1:**
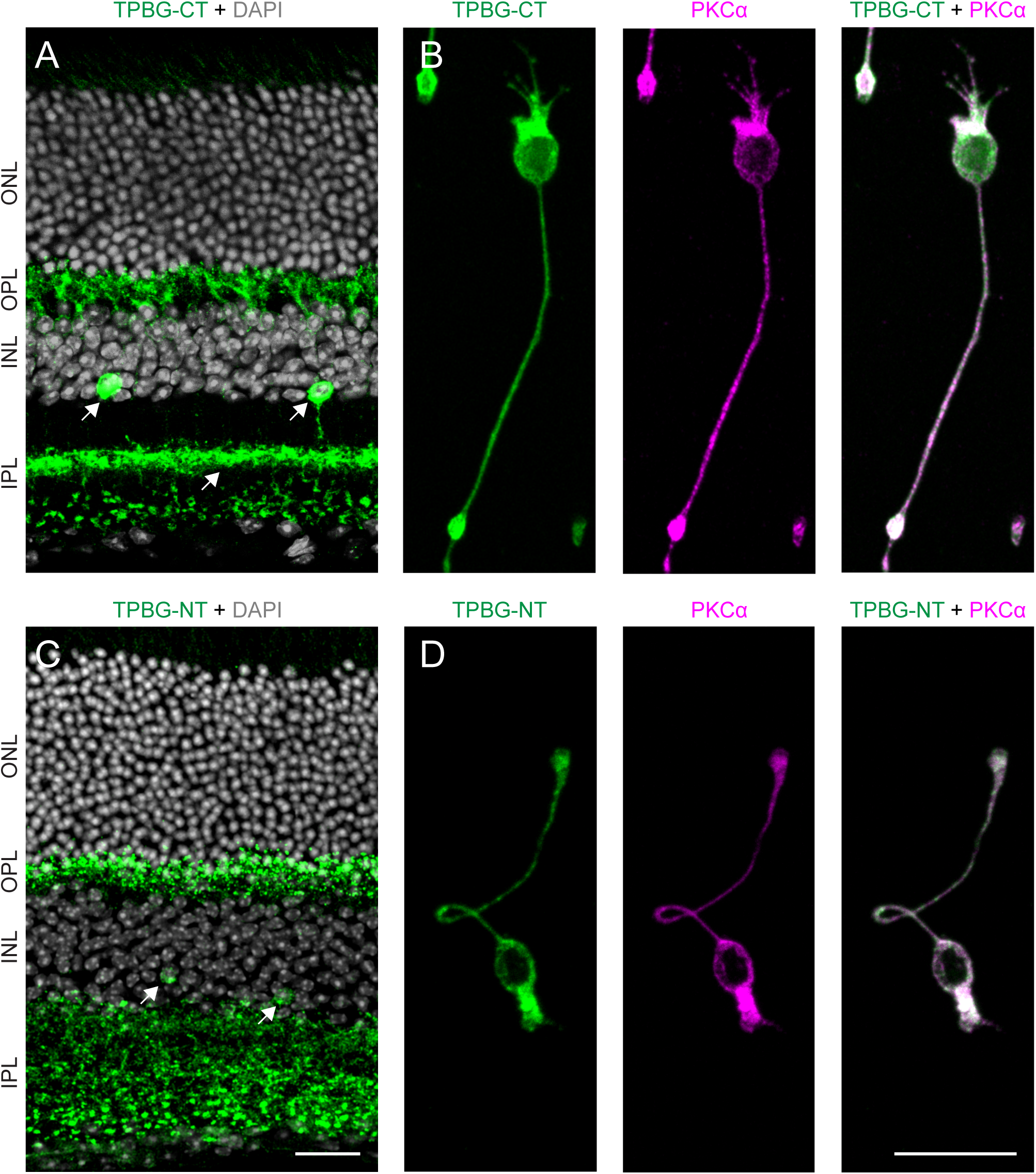
TPBG in the retina. Confocal microscopy of wild type adult retina sections and dissociated rod bipolar cells from wild type adult retina labeled with TPBG-CT (green A and B) or TPBG-NT (green, C and D). Double labeling with DAPI (grey, A and C) was used to highlight the layers of the neural retina and PKCα (magenta, B and D) was used to identify dissociated rod bipolar cells. Arrows point to TPBG-positive amacrine cell bodies in the INL and dendritic laminations in the IPL. Scale bars: 20 μm. ONL: outer nuclear layer; OPL: outer plexiform layer; INL: inner nuclear layer; IPL: inner plexiform layer.

Immunofluorescent double labeling of TPBG and PKCα confirms localization of TPBG to RBCs in retina sections (Figure 2). TPBG-CT immunofluorescence can be seen throughout the dendritic branches of RBCs in the OPL, as well as weaker labeling of their cell bodies in the outer INL (Figure 2A), and their synaptic terminals in the IPL (Figure 2B). In the OPL, anti-PKCα labels RBC cell bodies and dendrites, with the densest labeling occuring in the proximal dendrites just above the cell bodies. Anti-TPBG-CT, on the other hand, labels both the proximal and distal RBC dendrites. In the IPL, nearly complete co-localization of TPBG and PKCα in sublamina 5 confirms localization of TPBG to RBC synaptic terminals. Similar to Figure 1, a sparse population of amacrine cells was strongly reactive with anti-TPBG-CT (arrows), with cell bodies localizaed to the bottom of the INL and dendritic projections to a dense, narrow layer in the center of the IPL.

**Figure 2:**
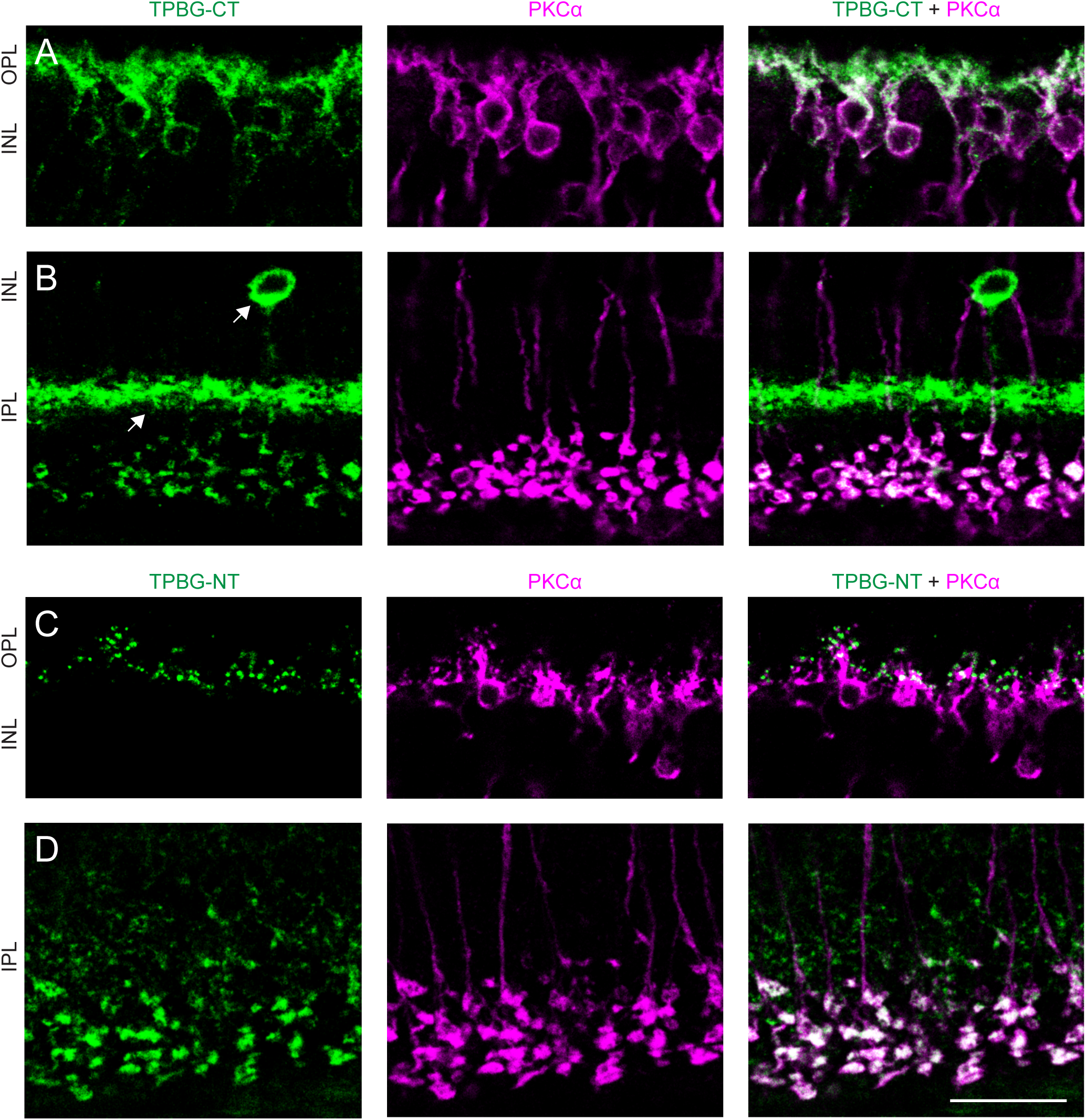
TPBG is expressed in rod bipolar cells. Confocal microscopy of wild type adult retina sections double labeled with TPBG-CT (A and B) or TPBG-NT (C and D) and an antibody against PKCα (magenta). Arrows point to TPBG-positive amacrine cells. Scale bar: 20 μm. OPL: outer plexiform layer; INL: inner nuclear layer; IPL: inner plexiform layer.

Immunofluorescent labeling with anti-TPBG-NT (Figure 2C and D) reveals punctate immunoreactivity in the OPL and widespread punctate labeling in the IPL that is not confined to sublamina 5. The TPBG-positive amacrine cell bodies and dendrites seen in Figure 1 and 2B are occasionally seen. In the OPL, there is much sparser but more punctate immunoreactivity (Figure 2C) than with anti-TPBG-CT. There is less overlap between anti-TPBG-NT and anti-PKCα, particularly in the RBC dendritic tips, which exhibit the strongest TPBG-NT labeling but are only weakly labeled for PKCα. In the IPL, there is punctate labeling across the entire width of the synaptic layer (Figure 2D), yet RBC synaptic terminals are still prominent, and all PKCα-positive RBC terminals are also labeled by anti-TPBG-NT.

### TPBG is localized to the plasma membrane and cytoplasmic compartments in transfected HEK293 cells

Based on its structure and similarities with other LRR proteins, TPBG is expected to be oriented in the plasma membrane with its N-terminal LRRs facing the extracellular space and its C-terminal tail in the cytoplasm. Since immunofluorescence in retina sections could not clearly reveal the subcellular distribution of TPBG in RBCs, and to better visualize its distribution within mammalian cells, HEK293 cells were transfected with full-length mouse TPBG and labeled with either the C-terminal or N-terminal TPBG antibodies (Figure 3). In fixed and permeabilized cells, both antibodies labeled cytoplasmic structures (Figure 3A). To prevent intracellular access, the antibodies were also added directly to live transfected cells (Figure 3B). Anti-TPBG-CT shows no immunoreactivity in the live cells, confirming the intracellular localization of the C-terminus. Anti-TPBG-NT, on the other hand, labeled the extracellular surface of live transfected HEK293 cells, confirming the predicted orientation of TPBG with the N-terminal, LRR domain facing out. Additionally, both antibodies labeled thin filopodia-like membrane projections (arrows in Figure 3B). Deconvolution of a Z-projection of anti-TPBG-CT immunofluorescence in a TPBG-transfected HEK293 cell (Figure 3C) revealed thin filopodia-like projections, suggesting plasma membrane localization of TPBG, and bright clusters of immunofluorescence that appear to be localized to the very tips of the projections (arrows in Figure 3C).

### TPBG is present in distal RBC dendrites in the OPL

To examine the dendritic distribution of TPBG in the mouse OPL, we compared TPBG-CT or TPBG-NT immunoreactivity with that of GPR179 and CtBP2 (Figure 4). GPR179 is localized exclusively to the dendritic tips of ON-bipolar cells, where it is responsible for anchoring the RGS7-Gβ5 complex to the postsynaptic membrane (Orhan et al., 2013; Orlandi, Cao, & Martemyanov, 2013; Tayou et al., 2016) in proximity to mGluR6 (Morgans et al., 2007; Ray et al., 2014). TPBG-CT-labeled puncta do not completely overlap GPR179 puncta at the very tips of the RBC dendrites, but do closely associate with them (3C, arrows). TPBG-NT puncta, on the other hand, strongly co-localize with GPR179 puncta in the OPL (3D, arrows). In the OPL, antibodies against CtBP2 label ribeye, a protein component of horseshoe-shaped synaptic ribbon structures in both cone and rod photoreceptor synaptic terminals (Schmitz et al., 2000). Proteins that localize to the tips of RBC dendrites can be visualized as puncta within the concavity of the ribeye horseshoe shape (Morgans et al., 2007). Both TPBG antibodies reveal punctate immunoreactivity within a subset of CtBP2-positive synaptic terminals (Figures 3E and F, arrows), consistent with the presence of TPBG near the tips of RBC dendrites.

**Figure 3:**
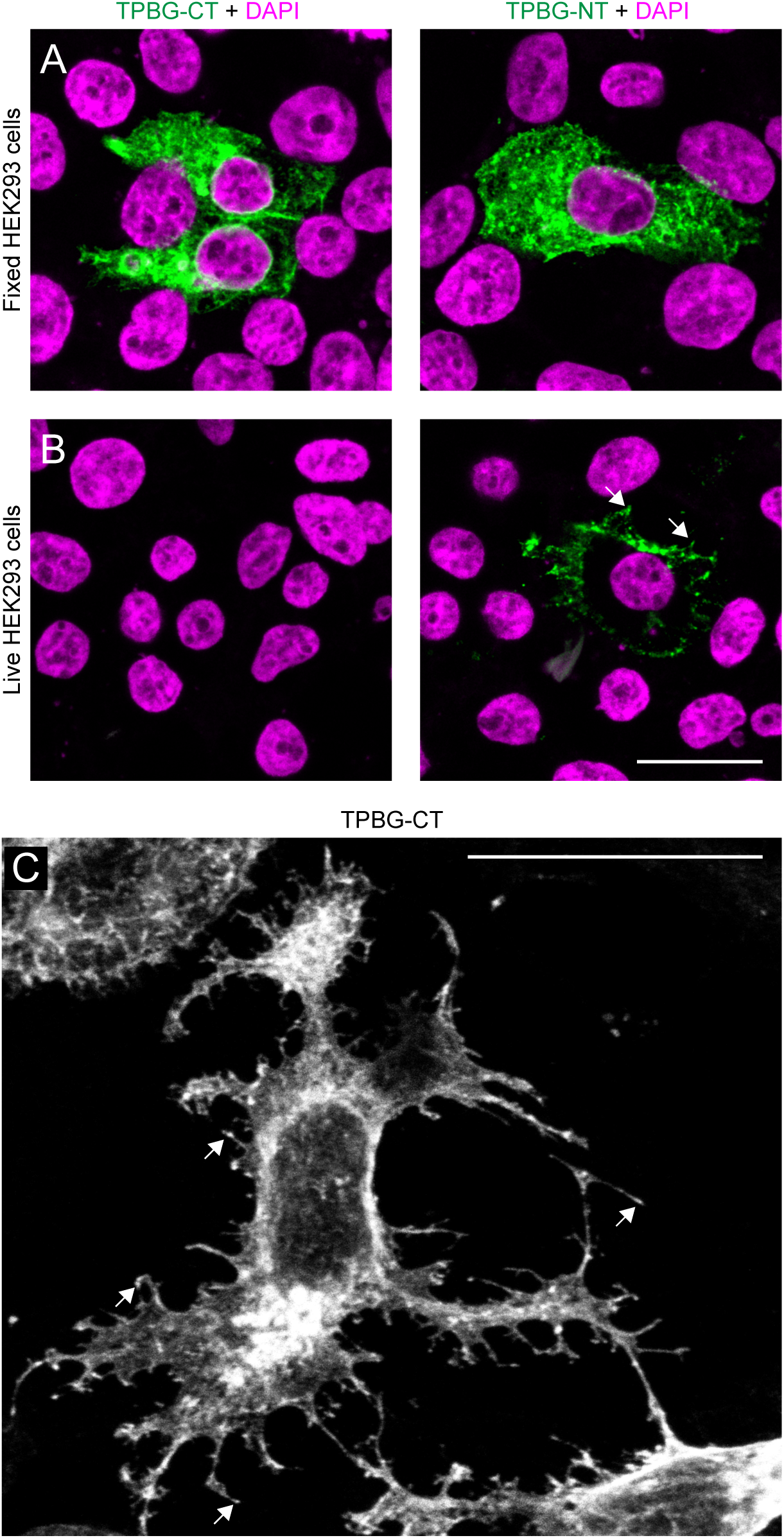
TPBG is localized to the extracellular surface and cytoplasmic compartments in transfected HEK293 cells. Confocal microscopy of HEK293 cells transfected with cDNA corresponding to full-length mouse TPBG inserted into the pCMV-sport6 vector. (A) Fixed and permeabilized cells were labeled with TPBG-CT and TPBG-NT. (B) Live cells were also labeled with TPBG-CT and TPBG-NT. (C) Z-projection of a TPBG-positive HEK293 cell labeled with TPBG-CT and processed using Huygens Hyvolution 2 deconvolution algorithms. Scale bars: 20 μm.

**Figure 4:**
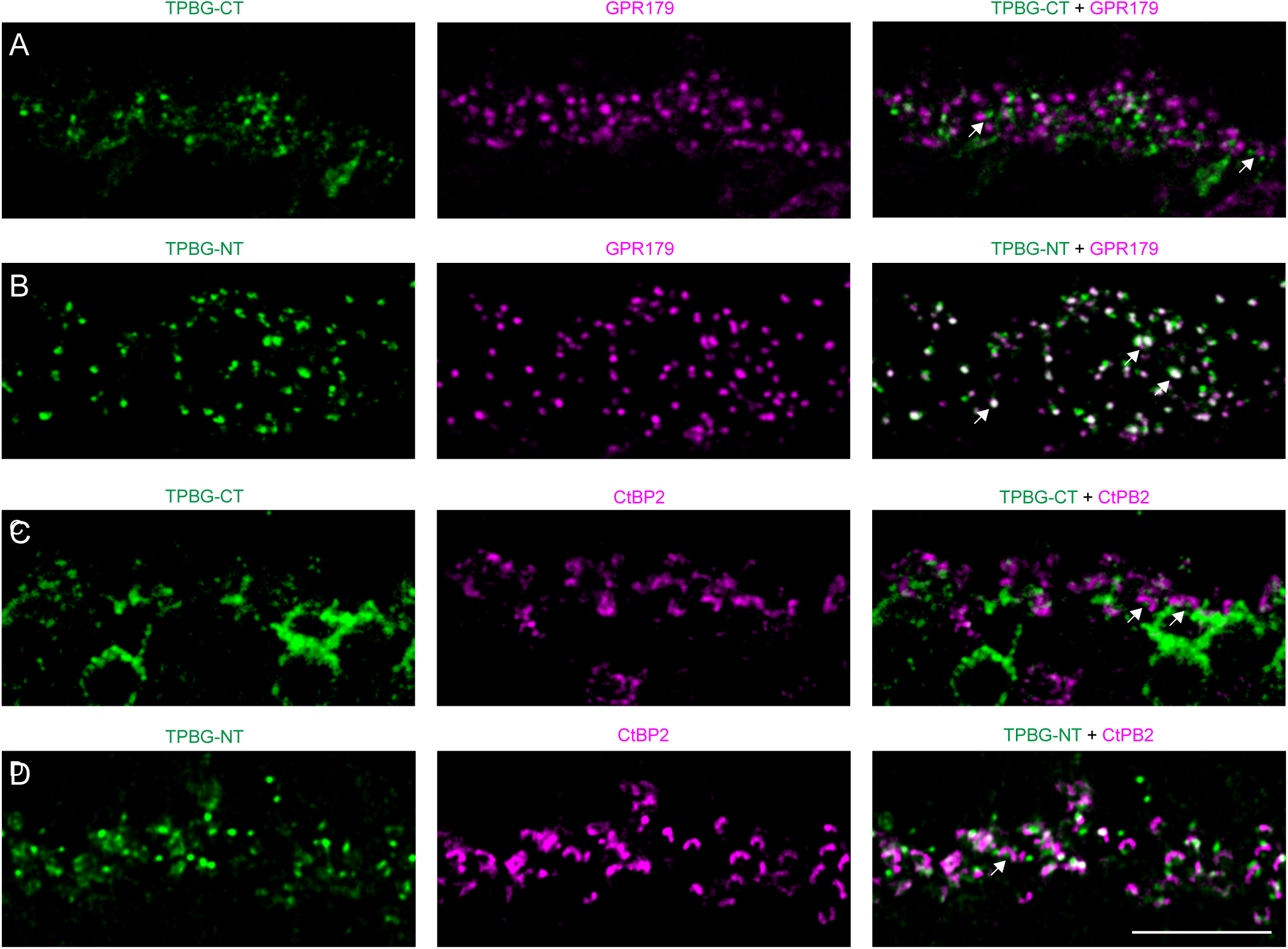
TPBG is present near RBC dendritic tips in the OPL. Confocal microscopy of wild type adult retina sections with TPBG-CT or TPBG-NT (green) and GPR179 (magenta, A and B) and CtBP2 (magenta, C and D). Scale bar: 10 μm.

### TPBG-positive amacrine cells

The IPL is typically divided into two layers corresponding to whether it receives input from ON or OFF bipolar cells, and can be further subdivided into five sublamina (2 OFF sublamina and 3 ON sublamina) based on labeling with immunofluorescence markers. One such IPL marker, calretinin, labels three distinct synaptic layers in the IPL corresponding the sublamina 1/2, 2/3, and 3/4 boundaries (Haverkamp & Wässle, 2000). The uncharacterized TPBG-positive amacrine cells project their dendrites to the middle of these three layers (Figure 5), overlapping with calretinin at the boundary between sublamina 2 and 3 between the ON and OFF regions of the IPL. This is similar to the IPL stratification of the TPBG-positive amacrine cell reported by Imamura et al. in 2006.

**Figure 5:**
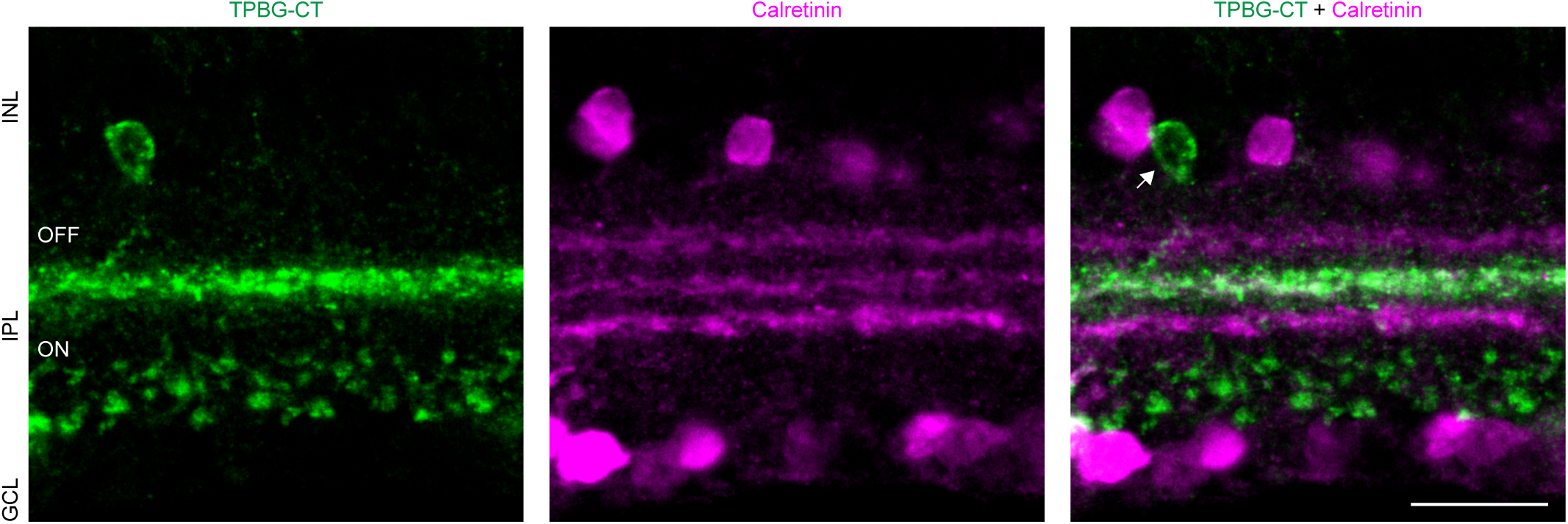
TPBG labels an uncharacterized population of amacrine cells. Confocal microscopy of wild type adult retina sections with TPBG-CT (green) and calretinin (magenta). Layers were determined by DAPI nuclear staining (not shown). INL: inner nuclear layer; IPL: inner plexiform layer; GCL: ganglion cell layer. Scale bar: 10 μm.

### TPBG immunofluorescence is altered under dark-adapted and knockout conditions

We looked for changes in TPBG expression in dark-adapted, PKCα knockout, and TRPM1 knockout retinas by immunofluorescent labeling and by immunoblotting retinal lysates. For the immunoblot analysis, bands at around 70 kDa corresponding to TPBG were normalized to β-actin bands within each sample to control for changes in total protein between samples. Dark-adaption resulted in a significant reduction in TPBG-CT immunoreactivity in both the OPL and the IPL, including the amacrine cell dendritic layer (Figure 6A), though surprisingly, no differences in TPBG levels were seen by immunoblot analysis of lysates from light-adapted and dark-adapted retinas (Figure 6B; light-adapted: 1.491, dark-adapted: 1.852, t-test p-value: 0.121, n=3 for both groups). PKCα knockout resulted in a slight reduction of TPBG-CT immunoreactivity in the OPL and in RBC terminals in the IPL, while the amacrine cell dendritic labeling was unchanged (Figure 6C). Total TPBG expression was unchanged between wild type and PKCα knockout retinal lysates when measured via immunoblot (Figure 6D; WT: 0.715, PKCα KO: 0.664, t-test p-value: 0.396, n=3 for both groups). In TRPM1 knockout retina, TPBG-CT immunoreactivity in RBCs was practically abolished when compared to wild type, but the amacrine cell labeling was unaffected (Figure 6E). Once again, there was no difference in total TPBG expression between wild type and knockout retina lysates as measured by immunoblot analysis(Figure 6F; WT: 0.698, TRPM1 KO: 0.748, t-test p-value: 0.710, n=3 for both groups). In contrast to TPBG-CT immunofluorescence, NT immunofluorescence and immunoblots were unchanged by any of the assessed conditions (not shown).

**Figure 6:**
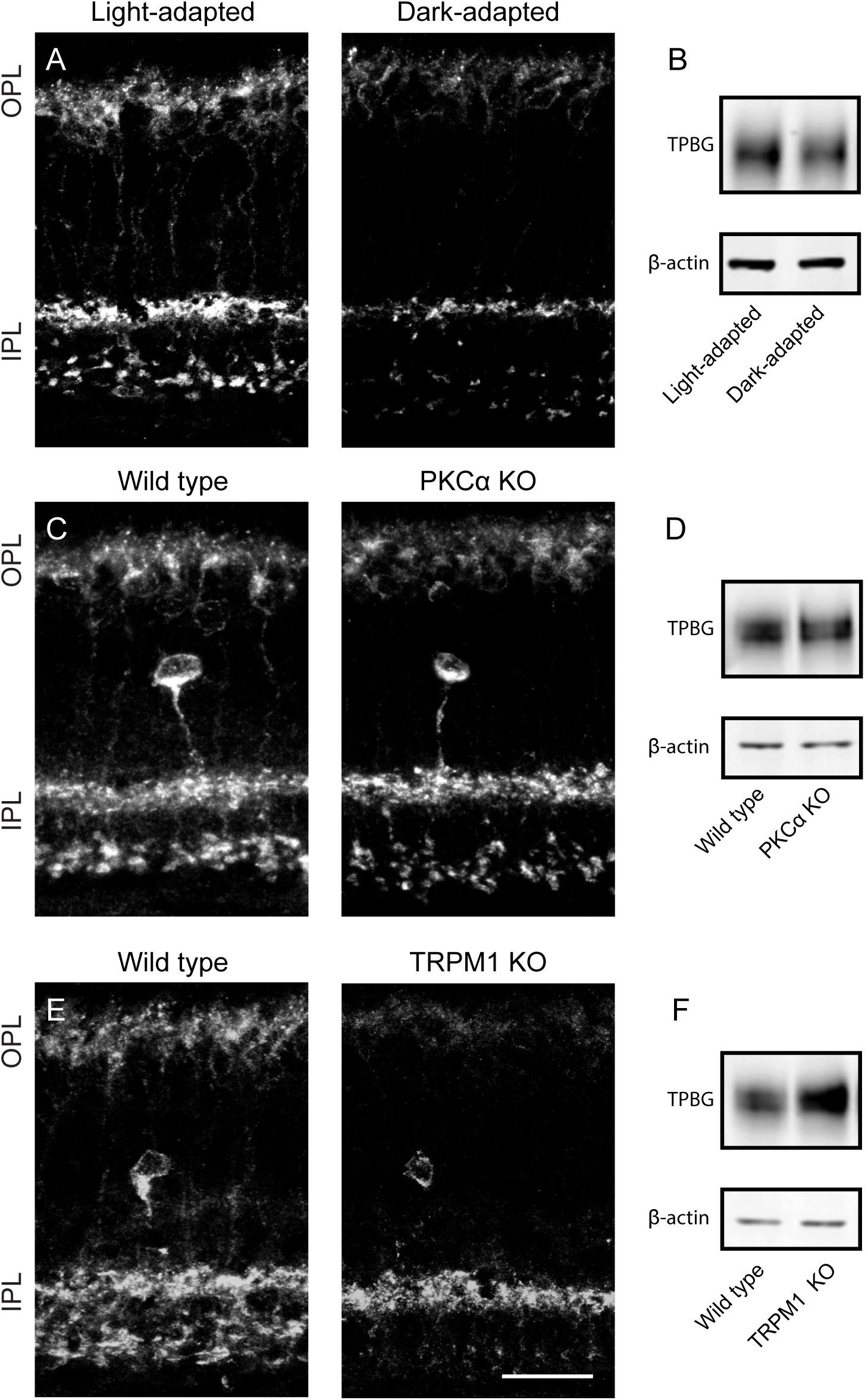
C-terminal TPBG immunofluorescence is altered under dark adapted and knockout conditions. Confocal microscopy of retina sections and immunoblots from (A) wild type light-adapted and (B) 12-hour dark-adapted mice, (C) wild type and (D) PKCα knockout mice, and (E) wild type and (F) TRPM1 knockout mice. There were no significant differences in immunoblot band intensity between any conditions after normalizing to β-actin (n=3 for all conditions).

### TPBG expression in the retina increases around eye opening

To analyze the time-course of TPBG expression in the developing mouse retina, retinal lysates were extracted at different postnatal (P) time-points and probed via immunoblot with antibodies to TRPM1 and either TPBG-CT or TPBG-NT (Figure 7A). Both TPBG antibodies detect a significant increase in retinal TPBG expression around eye opening, which occurred between P12 and P13, and is concomitant with an increase in the expression of TRPM1. The increase in TPBG expression around eye opening was also apparent in sections made from postnatal mouse retinas labeled with both TPBG antibodies. TPBG immunofluorescence was undetectable in the P0 retina (not shown). In the P6 retina, the TPBG-positive amacrine cell bodies and dendritic projections are just visible when labeled against TPBG-CT (Figure 7B, left, arrows). By P11, TPBG can be faintly seen in dendritic structures in the OPL (Figure 7C, arrows), but is still absent from RBC synaptic terminals in the IPL. At P12 (Figure 7D), TPBG immunofluorescence in the OPL has drastically increased, as has amacrine cell and RBC synaptic terminal labeling in the IPL. By P13 (Figure 7E), after eye opening, RBC dendrites in the OPL are clearly visible when labeled by TPBG-CT, while anti-TPBG-NT puncta have brightened significantly.

**Figure 7:**
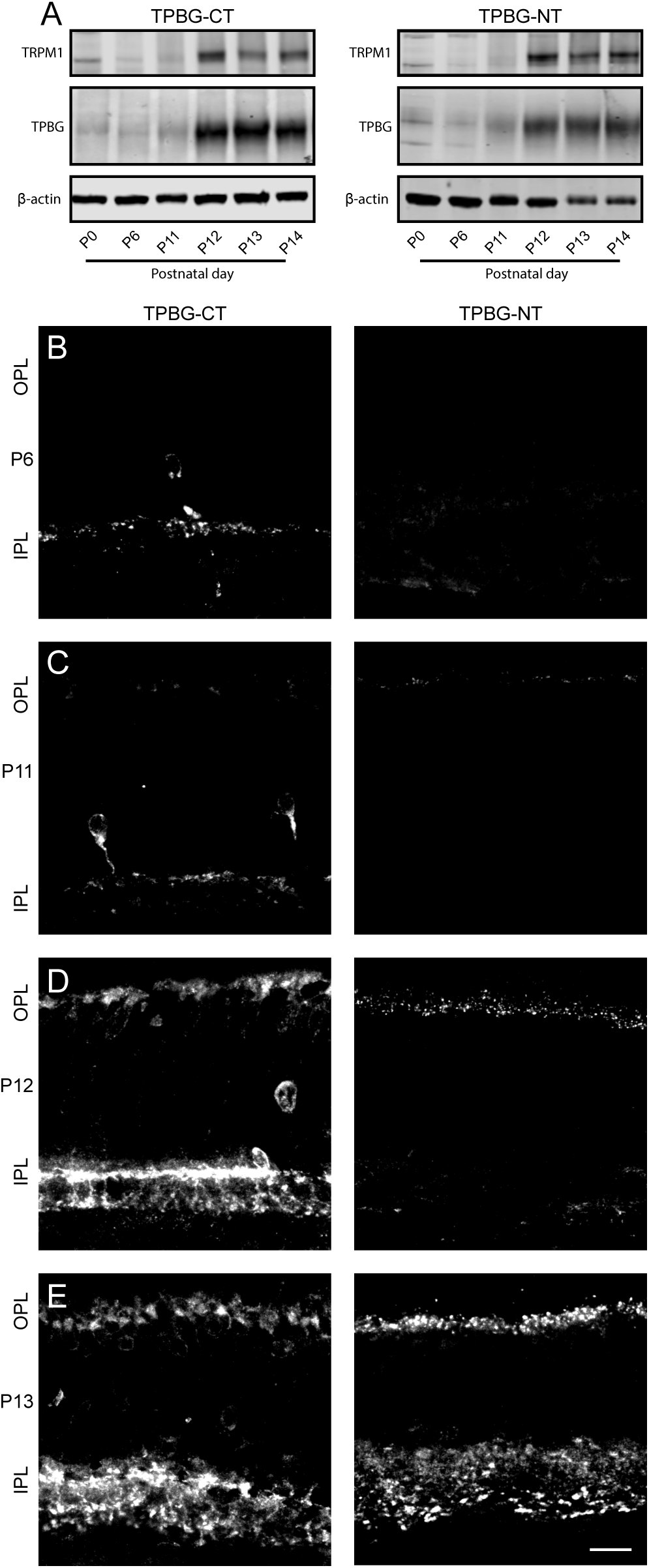
TPBG expression increases around eye opening. (A) Immunoblot of wild type retinal lysates prepared at different developmental time-points from the same litter and probed for TPBG-CT (left) or TPBG-NT (right), β-actin, and TRPM1. Confocal immunofluorescence microscopy of wild type mouse retina sections extracted at P6 (B), P11 (C), P12 (D), and P13 (E) and labeled for TPBG-CT (left) or TPBG-NT (right). Eye opening occurred between P12 and P13. Retinal layers were determined by DAPI nuclear staining. Scale bar: 20 μm.

## 4. Discussion

In this study, we have described the localization and expression patterns of TPBG in the mouse retina using two antibodies against intracellular and extracellular epitopes of TPBG. We found TPBG immunofluorescence primarily in the dendrites and synaptic terminals of RBCs and in the cell bodies and dendritic projections of an uncharacterized class of amacrine cell that stratifies in the middle of the IPL (Figures 1 and 2). In HEK293 cells, transfected TPBG was primarily localized to the cell membrane and in the tips of membrane projections (Figure 3). In the OPL, antibodies against both C-terminal and N-terminal epitopes label puncta closely associated with the tips of the RBC dendrites (Figure 4), co-localizing with or near GPR179 and CtBP2. In the IPL, both antibodies against TPBG label the membrane and cytoplasm of RBC synaptic terminals, overlapping with PKCα. Additionally, TPBG-NT appears to label the synaptic terminals of other cell types in the IPL, suggesting possible cross-reactivity with other leucine-rich repeat-containing proteins.

The N-terminal extracellular region of TPBG contains a heavily glycosylated (Shaw et al., 2002) leucine-rich repeat domain, a common site of protein-protein interactions (Zhao et al., 2014) and several similarly structured transmembrane proteins have recently been identified as vital for the development of the rod-RBC synapse or for localization of RBC synaptic transduction components. ELFN1, a synaptic adhesion LRR protein expressed in rod photoreceptors, forms trans-synaptic complexes with mGluR6 in RBC dendrites, and is required for the development of a functional synapse (Yan Cao et al., 2015; Dunn, Patil, Cao, Orlandi, & Martemyanov, 2018). Two other synaptic LRR proteins, LRIT3 and nyctalopin, are required for the localization of TRPM1 to the tips of ON bipolar cell synapses (Y Cao et al., 2011; Neuillé et al., 2017, 2015; Pearring et al., 2011). Our immunofluorescence data shows that TPBG is localized in distal RBC dendrites and in the RBC synaptic terminals, suggesting a possible role in the development or maintenance of the rod-RBC and RBC-AII synapses. LRR proteins in the IPL that could give rise to cross-reactivity with TPBG-NT include LRIT1 (Ueno et al., 2018), nyctalopin (Morgans, Ren, & Akileswaran, 2006), and synaptic adhesion proteins such as FLRTs (Visser et al., 2015), and neuroligins (M. Hoon et al., 2011; Mrinalini Hoon, Krishnamoorthy, Gollisch, Falkenburger, & Varoqueaux, 2017).

Immunofluorescent labeling with the C-terminal TPBG antibody was markedly affected by the activity state of the RBCs, with reduced immunofluorescence in dark-adapted, PKCα knockout, and TRPM1 knockout retinas (Figure 6). In contrast, N-terminal immunofluorescence was unaffected, and the total amount of TPBG detected on immunoblots with both TPBG antibodies was unchanged. It is possible that the variations in immunofluorescence observed with anti-TPBG-CT are due to differences in accessibility of the C-terminal epitope. The C-terminal intracellular region of TPBG is capped with a class 1 PDZ-interacting motif containing two serines (S422 and S424) that we have shown to be targets of PKCα-dependent phosphorylation (Wakeham et al., 2019). Phosphorylation of PDZ-interacting motifs has been shown to regulate binding to PDZ domains. For example, the C-terminal PDZ-interacting motifs of NMDA receptor subunits NR2A and NR2B, end with the amino acids SDV, like TPBG, and phosphorylation of the serine blocks binding of the receptor to the PDZ domain of PSD95 (Chung, Huang, Lau, & Huganir, 2004). It is not known what PDZ domain proteins may interact with the C-terminal, PDZ-interacting motif of TPBG in the retina; however, it has been demonstrated by yeast two-hybrid screening to bind to the PDZ domain of GIPC1, a scaffolding protein that regulates cell surface expression of GPCRs (Awan et al., 2002) that is expressed in RBCs (Shekhar et al., 2016). Together, our immunofluorescence and immunoblot results suggest that under conditions where TPBG-CT immunofluorescence is reduced (i.e. dark-adapted conditions, PKCα knockout and TRPM1 knockout), the C-terminus of TPBG may be bound to another protein, possibly containing a PDZ domain, rendering the C-terminal epitope inaccessible to anti-TPBG-CT. Phosphorylation of PDZ-interacting motifs typically abolishes binding to PDZ domains (Chung et al., 2004; Lee & Zheng, 2010), thus phosphorylation of the TPBG C-terminal serine residues in light-adapted, active RBCs may prevent binding of the interacting protein, permitting antibody binding to the C-terminal epitope.

In the developing mouse retina, TPBG expression is undetectable at birth. The presence of TPBG in amacrine cells is visible by P6, whereas RBC expression increases dramatically just prior to eye opening at P11-P12 (Figure 6). This pattern of increased expression approaching eye opening matches that of TRPM1, and is coincident with the gene expression patterns of many other proteins associated with the final stages of development of signal transduction machinery and establishment of bipolar cell morphology (Blackshaw et al., 2004; Dorrell, Aguilar, Weber, & Friedlander, 2004; Mu et al., 2001; Zhang, Kolodkin, Wong, & James, 2017). In embryonic development, TPBG alters Wnt and cytokine signaling to modulate cytoskeletal rearrangement and cell morphology (Kagermeier-Schenk et al., 2011; McGinn et al., 2012; Southgate et al., 2010). In the developing mouse retina, Wnt signaling between rods and RBCs is required for functional synaptic targeting and OPL lamination, and CRISPR-mediated inactivation of Wnt5 resulted in rod/RBC mistargeting and the formation of ectopic OPL (Sarin et al., 2018). Similarly, Ccl5-mediated chemokine signaling was found to be required for RBC axon terminal targeting to AII amacrine cell dendrites in the IPL (Duncan et al., 2018), suggesting that both Wnt- and chemokine-dependent mechanisms are active during RBC dendritic and axonal development. No relationship between TPBG and Wnt or chemokine signaling in the retina have yet been shown. However, as TPBG expression in RBCs increases concurrently with the activation of developmental processes dependent on these signaling pathways, it’s possible that TPBG is modulating similar signaling pathways in the retina RBCs.

We found that both antibodies against TPBG label an unidentified population of amacrine cell with a low-density distribution of cell bodies in the INL and dense dendritic stratification between the ON and OFF sublamina in the IPL (Figure 5). This TPBG-positive amacrine cell was first identified in a study focusing on TPBG in mouse olfactory bulb granule cells (Imamura et al., 2006). Based on similarities in cell body distribution and dendritic lamination density, we hypothesize that this might be the Type 47 (ac53-59) amacrine cell identified via connectomic reconstruction of the IPL, which was shown to receive inputs from cone bipolar cell 5D (XBC) and project outputs to two distinct populations of ganglion cells between the ON and OFF sublamina (Helmstaedter et al., 2013). The sparse distribution and dense lamination, combined with the connectome data from Helmstaedter et al., indicates that this TPBG-positive amacrine cell may have a wide dendritic field and be coupled to both the ON and OFF pathways. This is further supported by the fact that amacrine cell TPBG labeling was reduced in dark-adapted retina sections but not in PKCα or TRPM1 knockout. This could be explained by the fact that, while PKCα knockout only affects RBCs and TRPM1 knockout only affects the ON-pathway, dark-adaption suppresses activity of both ON and OFF pathways.

This study found that TPBG is localized to the dendrites and synaptic terminals of RBCs and the cell bodies and dendritic projections of an uncharacterized class of amacrine cell. C-terminal TPBG immunofluorescence was strongly dependent on the activity state of RBCs, possibly due to reduced epitope access caused by PDZ domain-containing binding partners. TPBG is expressed in both developing and adult retina, suggesting a continuing role in RBC cellular physiology before and after eye opening. Based on TPBG’s role in embryonic development, cancer tissue, and the olfactory bulb, and its structural similarities to other synaptic LRR proteins, it is reasonable to predict that TPBG may be involved in the development and maintenance of RBC synaptic morphology.

## Acknowledgments

The authors would like to thank Tammie L Haley for providing retina sections for immunofluorescent labeling. This work was supported by National Institutes of Health grants R01EY022369 and 5P30EY010572.

## Conflict of interest statement

The authors declare no conflicts of interest.

## Author contributions

study design: CMW, CWM; experimentation: CMW, GR; figure and manuscript preparation: CMW, CWM; editing and review: CMW, GR, CWM

## Abbreviations

CtBP2: C-terminal binding protein 2
GCL: ganglion cell layer
GPR179: G protein-coupled receptor 179
INL: inner nuclear layer
IPL: inner plexiform layer
LRR: leucine-rich repeat
OPL: outer plexiform layer
PKCα: protein kinase C-alpha
RBC: rod bipolar cell
TPBG: trophoblast glycoprotein
TRPM1: transient receptor potential cation channel subfamily M member 1

